# Single-domain antibodies efficiently neutralize SARS-CoV-2 variants of concern, including Omicron variant

**DOI:** 10.1101/2021.11.24.469842

**Authors:** Irina A. Favorskaya, Dmitry V. Shcheblyakov, Ilias B. Esmagambetov, Inna V. Dolzhikova, Irina A. Alekseeva, Anastasia I. Korobkova, Daria V. Voronina, Ekaterina I. Ryabova, Artem A. Derkaev, Anna V. Kovyrshina, Anna A. Iliukhina, Andrey G. Botikov, Olga L. Voronina, Daria A. Egorova, Olga V. Zubkova, Natalia N. Ryzhova, Ekaterina I. Aksenova, Marina S. Kunda, Denis Y. Logunov, Boris S. Naroditsky, Alexandr L. Gintsburg

## Abstract

Virus-neutralizing antibodies are one of the few treatment options for COVID-19. The evolution of SARS-CoV-2 virus has led to the emergence of virus variants with reduced sensitivity to some antibody-based therapies. The development of potent antibodies with a broad spectrum of neutralizing activity is urgently needed. Here we isolated a panel of single-domain antibodies that specifically bind to the receptor-binding domain of SARS-CoV-2 S glycoprotein. Three of the selected antibodies exhibiting most robust neutralization potency were used to generate dimeric molecules. We observed that these modifications resulted in up to a 200-fold increase in neutralizing activity. The most potent heterodimeric molecule efficiently neutralized each of SARS-CoV-2 variant of concern, including Alpha, Beta, Gamma, Delta and Omicron variants. This heterodimeric molecule could be a promising drug candidate for a treatment for COVID-19 caused by virus variants of concern.

## INTRODUCTION

The COVID-19 pandemic, caused by severe acute respiratory syndrome coronavirus 2 (SARS-CoV-2), has resulted in over 250 million of infections and over 5 million deaths worldwide (November 2021, WHO). These numbers are still rising and COVID-19 disease remains a great challenge to public health system. The development of safe and effective treatment together with vaccination is a highly important goal for scientists and health care professionals over the world.

One of the approaches to the development of an effective therapeutic agent is the isolation of monoclonal antibodies that efficiently neutralize SARS-CoV-2 virus. Currently, several monoclonal antibodies already received emergency use authorization for COVID-19 treatment and post-exposure prophylaxis. These antibodies are bamlanivimab plus etesevimab, casirivimab plus imdevimab, sotrovimab and regdanvimab (1–4).

Along with conventional antibodies, camelid single-domain antibodies (also called nanobodies or VHHs) are promising candidates for the development of antibody-based therapies (5, 6). Camelids have unique heavy-chain antibodies that are devoid of light chains (7). Nanobodies are the minimal antigen-binding domains of these heavy-chain antibodies. They have several advantages, including the ability to recognize epitopes that are not accessible to conventional antibodies, increased stability of nanobodies, simplicity of generation of multivalent forms and low cost bacterial production (8). Currently, several nanobodies to SARS-COV-2 have been isolated showing neutralizing activity in *in vitro* and *in vivo* studies, which confirms their promising potential as therapeutic agents (9–15).

The evolution of SARS-CoV-2 virus has resulted in emergence of virus variants that have become more transmissible and less sensitive to neutralizing antibodies. The spread of these new virus variants has reduced the efficacy of vaccines and some therapeutic antibodies. The list of these variants of concern (VOCs) consist of B.1.1.7 (Alpha), B.1.351 (Beta), B.1.1.28/P.1 (Gamma), B.1.617.2 (Delta) and B.1.1.529 (Omicron) variants (December 2021, WHO). The reduction of neutralization by antibodies is caused by mutations in S glycoprotein, including K417N/T, L452R, T478K, E484K and N501Y substitutions (16, 17). Recently appeared Omicron variant has more than 30 mutations in S glycoprotein that provide a considerable escape from neutralization by antibodies (18, 19). In this regard, it became necessary to isolate antibodies, the neutralizing activity of which will not be affected due to observed mutations, therefore, these antibodies or their cocktails will retain activity against each of VOCs.

Here we identified and characterized a panel of single-domain antibodies isolated from immune VHH library that specifically bind RBD of S glycoprotein. We assessed the neutralizing activity of the isolated antibodies in a microneutralization assay with live SARS-CoV-2 and selected three most potent antibodies. To increase the therapeutic potential, these clones were modified to homodimeric and heterodimeric forms, and the neutralizing activity against SARS-CoV-2 VOCs was investigated. The most potent heterodimeric form, P2C5-P5F8, exhibited activity against all tested virus variants at low concentration. These results indicate that P2C5-P5F8 heterodimer is a promising candidate for further research to develop COVID-19 antibody-based therapy.

## RESULTS

To identify SARS-CoV-2 neutralizing single-domain antibodies we immunized one Bactrian camel with SARS-CoV-2 RBD. The recombinant RBD protein was previously produced in CHO-S cells and purified. Immunization was performed using five sequential injections (Fig 1A). Blood was collected 5 days after final immunization and the serum RBD-specific antibodies titer was detected by ELISA. Post-immunization serum demonstrated potent binding to SARS-CoV-2 RBD with titer more than 1/200 000 (Fig 1B). The neutralizing activity of antibodies in the immunized serum was measured by the microneutralization assay using live SARS-CoV-2 B.1.1.1, the neutralizing antibodies titer was 1/1280.

**Figure 1.**
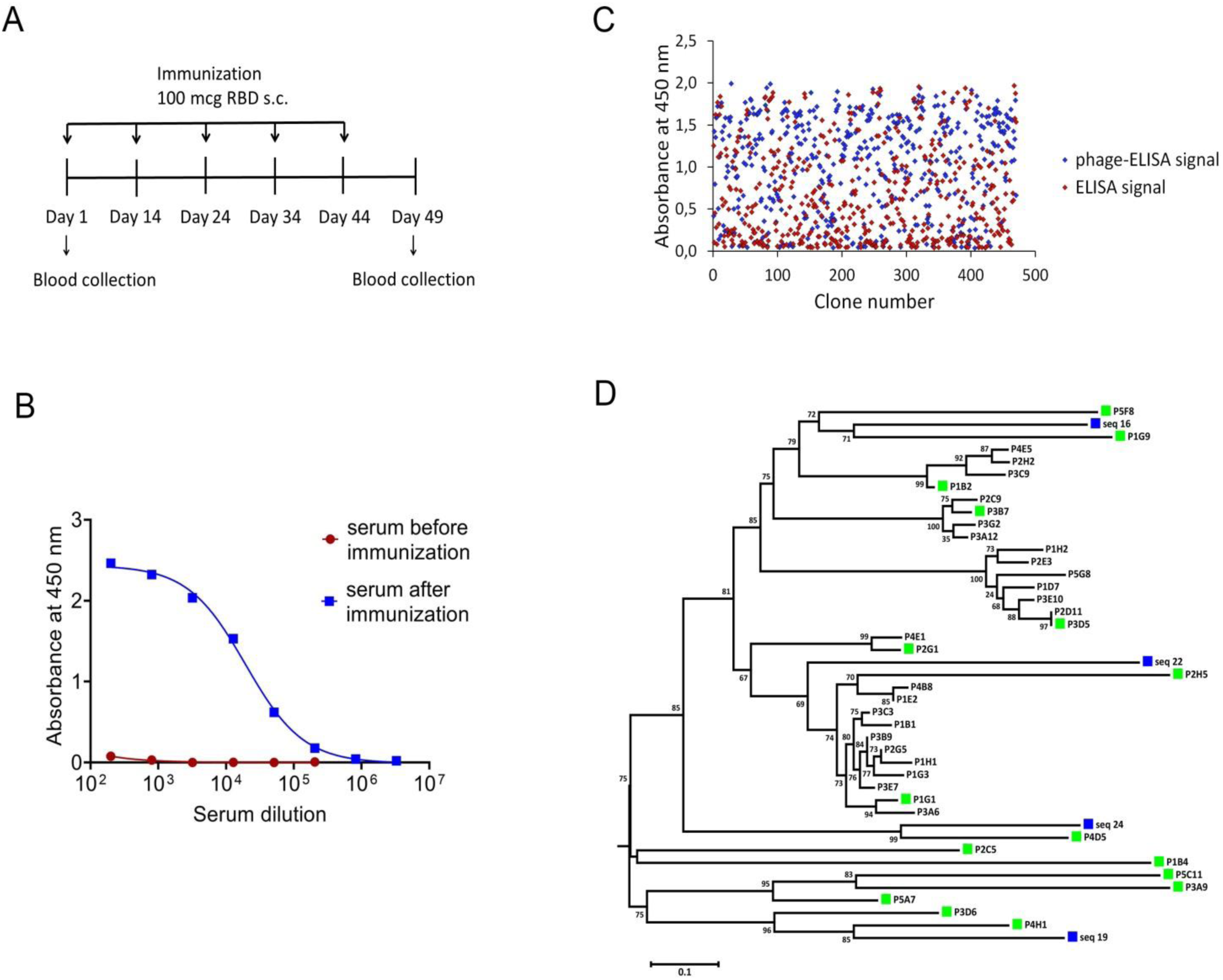
Isolation of RBD-specific nanobodies. (**A)** Immunization schedule. Bactrian camel was immunized with 100 mcg RBD subcutaneously (with complete Freund’s adjuvant), followed by four consecutive immunization with 100 mcg RBD subcutaneously (with incomplete Freund’s adjuvant). Blood samples were collected before immunization and five days after the last immunization. (**B)** RBD-specific antibodies in camel serum before and after immunization, detected by ELISA. The assay reveals a strong positive RBD-specific serological activity 5 days after the last immunization. (**C)** ELISA-based RBD-binders screening. A total of 212 individual clones with a strong positive ELISA signal were selected for sequencing. (**D)** Phylogenetic tree showing sequence diversity of 39 unique VHH clones from this study and four previously described naive single-domain antibodies of C. bactrianus (45), blue squares - naive single-domain antibodies of C. bactrianus, green squares – the clones selected for further analysis.

To isolate monoclonal RBD-specific nanobodies, we constructed a VHH phage display library by cloning VHH sequences from B cells of immunized camel into a phagmid vector. We performed one round of phage display followed by ELISA-based screening of individual clones to identify RBD-specific binders.

We used monoclonal phage-ELISA together with monoclonal ELISA to identify RBD binders with high expression levels and high solubility. A total of 212 clones showing positive phage-ELISA and ELISA signals were selected for sequencing (Fig 1C), which lead to the identification of 39 unique clones of nanobodies. After sequence alignment and phylogenetic analysis we selected 16 antibodies with highly diverse sequences for further research (Fig 1D). These nanobodies were expressed and purified, and the RBD binding activity of each antibody was analyzed by ELISA. We confirmed the binding of 14 clones with ELISA EC50 values ranged from 1.1 to 313.3 nM (Fig 2A). Most of the antibodies (13 out of 14) displayed strong positive signals with EC50 below 22 nM.

**Figure 2.**
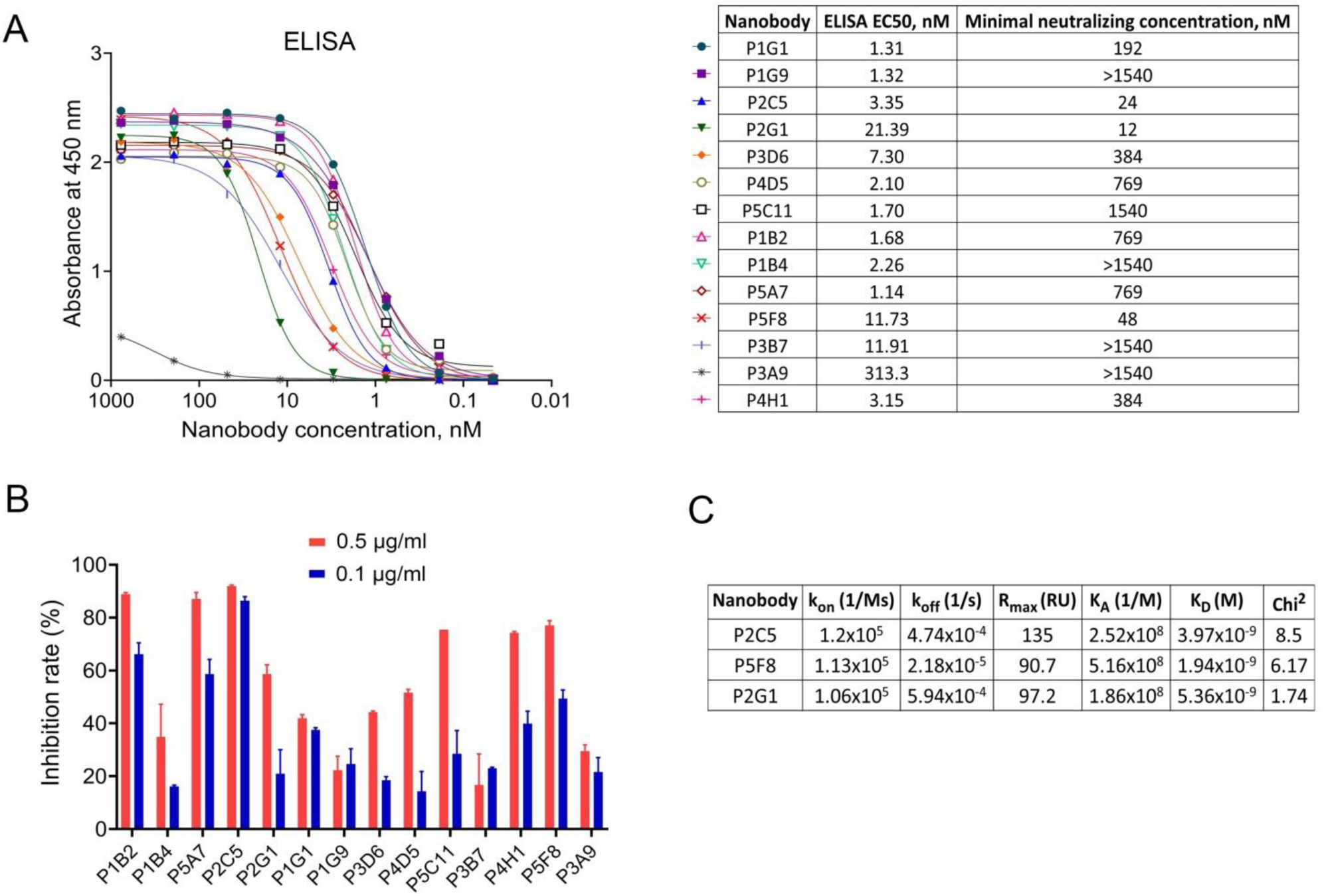
Characterization of the selected nanobodies. (**A)** RBD-binding activity of nanobodies by ELISA and neutralization activity of nanobodies by microneutralization assay using live SARS-CoV-2 virus. The minimal neutralizing concentration was defined as the lowest antibody concentration (highest antibody dilution) that completely inhibited the cytopathic effect of the virus in two or three from the three replicable wells. (**B)** Blocking of ACE2-RBD interaction measured by competitive ELISA. (**C**) Kinetic parameters of nanobodies interaction with RBD by SPR. Association (k_on_), dissociation (k_off_), maximal analyte binding capacity (R_max_), equilibrium association constants (K_A_), equilibrium dissociation constants (K_D_) and Chi^2^ for VHHs binding to RBD.

To examine the capacity of selected RBD binders to neutralize live SARS-CoV-2 virus we performed a microneutralization assay with inhibition of the cytopathic effect of virus as a marker of neutralization. We found that 10 of 14 nanobodies efficiently neutralized SARS-CoV-2 virus (B.1.1.1) at concentrations ranging from 12 nM to 1540 nM (Fig 2A). The three most potent antibodies were P2C5, P2G1 and P5F8, which completely inhibited the cytopathic effect of the virus at 24 nM, 12 nM and 48 nM, respectively.

We investigated the binding kinetics of P2C5, P2G1 and P5F8 neutralizing nanobodies via surface plasmon resonance (SPR). All antibodies tested bound RBD with high affinity, dissociation constants (K_D_) were 3.97 nM, 5.36 nM and 1.94 nM for P2C5, P2G1 and P5F8, respectively (Fig 2C).

Since the capacity of an antibody to neutralize SARS-CoV-2 virus is frequently associated with blocking the interaction between S glycoprotein and ACE2 receptor, we analyzed the ability of isolated nanobodies to block the RBD binding to ACE2 by competitive ELISA (Fig 2B). We revealed a decrease in ELISA signal for most of neutralizing nanobodies indicating antibodies competition with ACE2 for RBD binding. We observed that one of the most potent neutralizing antibodies, P2C5, blocked the ACE2-RBD interaction with an inhibition rate greater than 90% at 0.5 μg/ml.

In order to improve the antiviral property of antibodies we produced homodimer and heterodimer forms of P2C5, P2G1 and P5F8 clones. Monomers were fused using a flexible GS-linker (Gly4Ser)_4_ for dimerization. The binding of generated dimeric molecules to the recombinant RBD protein was confirmed by ELISA. Analysis of the neutralization capacity in the microneutralization assay revealed a pronounced improvement in the neutralizing activity of several dimeric forms (Table 1). Notable, P2C5 homodimer inhibited cytopathic effect of the virus at 89 pM, thus the homodimer activity increased more than 200-fold compared to the monomer activity. We also observed that P2C5-P5F8 heterodimer form was up to 100 times more active than the monomeric forms, this molecule exhibited neutralization activity at 178 pM.

**Table 1.** Neutralizing activity of the most potent monomers of nanobodies, homodimeric and heterodimeric forms of nanobodies in a microneutralization assay against live SARS-CoV-2 variants of concern.

| Antibody |  | Minimal neutralizing concentration of antibody, nM |  |  |  |  |  |
| --- | --- | --- | --- | --- | --- | --- | --- |
|  |  | B.1.1.1 | Alpha<br>(B.1.1.7) | Beta<br>(B.1.351) | Gamma<br>(B.1.1.28/P.1) | Delta<br>(B.1.617.2) | Omicron<br>(B.1.1.529) |
| Monomers | P2C5 | 24.04 | 48.08 | 96.15 | 48.08 | >1500 | 24.04 |
|  | P2G1 | 12.02 | 48.08 | 96.15 | 48.08 | 24.04 | >1500 |
|  | P5F8 | 48.08 | 48.08 | 96.15 | 96.15 | 48.08 | >1500 |
| Homodimers | (P2C5)2 | 0.089 | 0.178 | 0.356 | 0.356 | >1500 | 0.089 |
|  | (P2G1)2 | 11.36 | 22.73 | 45.45 | 45.45 | 45.45 | >1500 |
|  | (P5F8)2 | 5.68 | 5.68 | 22.72 | 22.72 | 11.36 | >1500 |
| Heterodimers | P2C5-P2G1 | 0.709 | 0.356 | 2.84 | 2.84 | 11.36 | 0.709 |
|  | P2C5-P5F8 | 0.178 | 0.089 | 0.356 | 0.356 | 2.85 | 0.709 |
|  | P2G1-P2C5 | 2.84 | 2.84 | 11.36 | 5.68 | 45.45 | 2.84 |
|  | P2G1-P5F8 | 45.45 | 45.45 | 181.82 | 45.45 | 90.91 | >1500 |
|  | P5F8-P2C5 | 5.68 | 11.36 | 22.73 | 22.72 | 90.91 | 5.68 |
|  | P5F8-P2G1 | 11.36 | 22.72 | 90.91 | 22.72 | 22.73 | >1500 |

With the emergence and spread of new variants of SARS-CoV-2 with increased transmissibility and reduced neutralization by antibodies it became necessary to isolate antibodies with a broad spectrum of neutralizing activity. We assessed the neutralizing activity of isolated antibodies against variants of concern (VOCs), including B.1.1.7 (Alpha), B.1.351 (Beta), B.1.1.28/P.1 (Gamma), B.1.617.2 (Delta) and B.1.1.529 (Omicron) variants (Table 1). P2C5 monomer neutralized B.1.1.7, B.1.351, B.1.1.28/P.1 and B.1.1.529 but was unable to neutralize B.1.617.2. P5F8 and P2G1 monomeric forms remained active against all tested VOCs (inhibited CPE at 12-96 nM) except Omicron variant. P5F8 monomer showed a 2-fold decrease in activity against B.1.351 and B.1.1.28/P.1. P2G1 showed a 2-4-fold decrease against VOCs. The most potent heterodimeric molecule P2C5-P5F8 exhibited activity against all VOCs neutralizing B.1.1.7, B.1.351, B.1.1.28/P.1, B.1.617.2 and B.1.1.529 variants at 89 pM, 356 pM, 356 pM, 2.85 nM and 709 pM, respectively. The most active P2C5 homodimer neutralized each of VOC except B.1.617.2 at a concentration 356 pM and lower, depending on the virus variant. Overall, P2C5-P5F8 heterodimer showed neutralizing activity against each of the tested VOCs and was a promising candidate for further investigations as a potential therapeutic agent.

## DISCUSSION

The SARS-CoV-2 virus gave rise to ongoing pandemic, which continues to claim lives and poses a threat to public health. Along with vaccination to prevent COVID-19, therapeutic antiviral agents are urgently needed to treat the infection.

One of the promising approaches for the viral infection treatment is an antibody-based therapy. Antibodies administration approved for immunoprophylaxis and treatment of viral infections caused by respiratory syncytial virus and ebolavirus (20, 21). Today, several monoclonal antibodies are already in use for COVID-19 treatment (bamlanivimab plus etesevimab, casirivimab plus imdevimab and sotrovimab). It was shown that the administration of these antibodies to nonhospitalized patients with symptomatic COVID-19 and risk factors for disease progression reduced the risk of hospitalization and death (1–4).

In addition to conventional antibodies, single-domain antibodies or nanobodies from camelid heavy-chain antibodies are being actively investigated as a potential therapeutic agent with a number of advantages. The small size and single-domain nature of nanobodies allow them to bind epitopes that are not available for conventional antibodies, in particular concave epitopes such as grooves and clefts (22–24). Nanobodies show strong binding affinities and also are highly stable across wide range of temperatures and pH (25). The production of nanobodies in bacteria is less expensive than classical antibody production. All this allows the use of single-domain antibodies for many of applications, including research, diagnostics and therapy (6, 8).

The main mechanism of virus-neutralizing activity of antibodies is associated with the prevention of viral entry into host cells. The entry of SARS-CoV-2 virus occurs after the interaction of S glycoprotein with cell receptor angiotensin-converting enzyme 2 (ACE2). Therefore, S glycoprotein, in particular its receptor-binding domain (RBD), is an attractive target for the development of neutralizing antibodies.

In our work, we immunized a Bactrian camel with the recombinant RBD protein for following isolation of effective neutralizing antibodies against SARS-CoV-2 virus. We generated a VHH phage display library and selected a panel of RBD-specific nanobodies. Using an *in vitro* microneutralization assay we identified three clones (P2C5, P5F8 and P2G1) of the most potent antibodies that completely inhibited the cytopathic effect of SARS-CoV-2 virus at minimal concentrations from 12 nM to 48 nM.

At the next stage, to enhance the neutralization potential of isolated nanobodies we generated homodimeric and heterodimeric forms of the most active antibodies. It was previously shown that multivalent forms of nanobodies are characterized by increased functionality by improving antigen-binding through avidity effect (26). For example, the dimeric form of anti-TNF VHH was up to 500 times more active than the monovalent form in rheumatoid arthritis model *in vivo* (27), homotrimeric molecule of anti-SARS-CoV-2 nanobody Nb21_3_ were 30-fold more potent in pseudovirus neutralization (14), dimeric forms of nanobodies to botulinum neurotoxin showed a higher protective activity *in vivo* (28). In our study, we revealed a 200-fold improvement of neutralization potential after dimerization of P2C5 nanobody. The bivalent molecule effectively neutralized SARS-CoV-2 virus at 89 pM. We also observed a pronounced increase in neutralizing activity in the case of P2C5-P5F8 heterodimeric form, which inhibited SARS-CoV-2 virus at 178 pM. In the case of the bivalent form of P5F8 we observed only a slight increase in activity. We assume that this may be due to the monovalent binding of P5F8 dimer to the S glycoprotein, for example as a result of the insufficient linker length to span the distance between epitopes in the trimeric S glycoprotein.

The emergence of SARS-CoV-2 variants with mutations in their RBD epitopes leads to the viral escape from neutralizing antibodies. It was shown that the virus-neutralization activity of sera samples obtained from individuals vaccinated with Pfizer/BNT162b2, Moderna/mRNA-1273 or Sputnik V decreased in the case of the B.1.351, B.1.1.28/P.1 and B.1.617.2 variants compared to wild-type virus (29–32). The most pronounced decrease in the virus-neutralization activity of sera is observed in the case of the B.1.1.529 (Omicron) variant (33-35). Escape mutations are also a challenge for antibody-based therapy development. Some therapeutic antibodies have become ineffective against VOCs, for example, the activity of bamlanivimab against Delta is reduced, the activity of etesevimab and casivimab against Beta variant is reduced (36). The activity of most neutralizing antibodies is escaped by the Omicron variant. Several monoclonal antibodies have completely lost their inhibitory activity against Omicron, including bamlanivimab, etesevimab, casirivimab, imdevimab and regdanvimab (18, 37). The activity of some previously isolated nanobodies also altered by escape mutations. The Beta strain drastically reduced the efficacy of the high-affinity nanobodies Nb20 and Nb21 (38). Thereby, it is urgently needed to identify nanobodies or their combinations that retain activity against SARS-CoV-2 variants of concern.

We investigated the activity of isolated potent nanobodies and their dimeric forms using a microneutralization assay against live SARS-CoV-2 VOCs. We found that P2C5-P5F8 heterodimer effectively neutralized all tested virus variants at low concentration, including Delta and Omicron VOCs. In particular it was observed that this dimeric molecule completely inhibited the cytopathic effect of the Alpha, Beta, Gamma, Delta and Omicron variants at concentrations 89 pM, 356 pM, 356 pM, 2.85 nM and 709 pM, respectively.

For the improvement of antiviral efficacy of obtained P2C5-P5F8 molecule further modifications may include the fusion of the heterodimer to human IgG1 Fc domain. Fc modifications are often used to increase serum half-life of nanobodies, as well as to involve Fc-mediated effector functions (antibody-dependent cell-mediated cytotoxicity, complement-dependent cytotoxicity, etc.) (28, 39).

Overall, we produced heterodimeric form of nanobodies that possess a broad spectrum of activity, overcoming escape mutations in VOCs. This heterodimeric molecule is a promising candidate for the further development of an effective antiviral agent for COVID-19 treatment especially for patients with risk factors for progression to severe COVID-19.

## MATERIALS AND METHODS

### Cell lines and viruses

CHO-S cells were obtained from Thermo Fisher Scientific (USA), cat. no. R80007. Vero E6 cells (ATCC CRL 1586) was obtained from Russian collection of vertebrate cell lines.

SARS-CoV-2 strains B.1.1.1 (hCoV-19/Russia/Moscow_PMVL-1/2020), B.1.351 (hCoV-19/Russia/SPE-RII-27029S/2021), B.1.617.2 (hCoV-19/Russia/SPE-RII-32758S/2021) and B.1.1.529 (hCoV-19/Russia/MOW-Moscow_PMVL-O16/2021) were isolated from nasopharyngeal swabs.

SARS-CoV-2 strains B.1.1.7 (hCoV-19/Netherlands/NoordHolland_20432/2020) and B.1.1.28/P.1 (hCoV-19/Netherlands/NoordHolland_10915/2021) were obtained from European Virus Archive Global.

### Cloning, expression and purification of recombinant SARS-CoV-2 RBD protein

The RBD nucleotide sequence of SARS-CoV-2 Wuhan-Hu-1 isolate (Genbank accession number MN908947, from 319 to 545 aa) was synthesized (Evrogen, Russia) and cloned into the pCEP4 mammalian expression vector (Thermo Fisher Scientific, USA). The signal peptide of alkaline phosphatase SEAP was inserted before RBD coding sequence, the His tag was fused to the end of the sequence. The obtained recombinant vector was transfected into CHO-S cells using CHOgro High Yield Expression System (Mirus Bio, USA). The culture supernatant containing the RBD protein was harvested after 10 days, the recombinant RBD protein was purified by IMAC using an AKTA start system (Cytiva, USA) and a HisTrap column (Cytiva, USA). The protein expression and purity was confirmed with SDS-PAGE.

### Bactrian camel immunization

Bactrian camel was immunized with recombinant SARS-CoV-2 RBD protein at a dose of 100 μg with complete Freund’s adjuvant subcutaneously followed by four booster immunizations with 100 μg RBD with incomplete Freund’s adjuvant. The interval between the first and second injections was 14 days; the intervals between all subsequent injections were 10 days. Before immunization, a small blood sample (5 ml) was collected as a control. 50 ml blood was collected 5 days after the last immunization and was used for peripheral blood mononuclear cells (PBMC) isolation and VHH phage display library construction.

### Phage display library construction

The library construction was performed as described earlier (28). Briefly, total RNA was isolated from PBMCs and then reverse transcribed to cDNA. VHHs coding sequences were PCR amplified and cloned into a pHEN1 phagmid vector (40). Recombinant phagmids were electroporated into TG1 cells (Lucigen, USA). A VHH library of 7.3 × 10^6^ individual clones was obtained.

### Phage preparation and biopanning

Phage preparation was performed according to the previous description (28). To select RBD-specific nanobodies one round of biopanning was performed. 5 μg of recombinant SARS-CoV-2 RBD protein were immobilized in the well of microtiter plate. After rinsing three times with PBS with 0.1% Tween 20 (TPBS), the well was blocked with blocking buffer (5% non-fat dried milk in TPBS) at 37 °C for 1 h. A total of ∼10^11^ phage particles in blocking buffer were added to the well and the plate was incubated at 37 °C for 1 h. Unbound phages were removed by washing with TPBS. The bound phages were eluted by trypsin (1 mg/ml). The eluted phage particles were used for infection of TG1 cells.

### ELISA

Screening for RBD-specific binders was performed using monoclonal phage ELISA and monoclonal ELISA. For monoclonal phage ELISA, individual clones were inoculated in 1 ml of 2xYT (Sigma, USA) containing 100 μg/ml ampicillin in 96-deep-well plates. Bacterial cells were infected with KM13 helper phage and incubated overnight at 30 °C. The plates were centrifuged, 50 μl of the supernatant were used for phage ELISA. For monoclonal ELISA, individual clones were cultured in 96-deep-well plates at 30 °C overnight. The next day, the cells were lysed by the freeze-thaw method, and then the plates were centrifuged. 50 μl of the supernatant were used for monoclonal ELISA. Immunoplates (Nunc MaxiSorp, Thermo Fisher Scientific, USA) were coated with recombinant RBD protein (100 ng per well) at 4 °C overnight. The plates were rinsed three times with TPBS and the wells were blocked with blocking buffer at 37 °C for 1 h. 50 μl of bacterial supernatants containing monoclonal recombinant phages (monoclonal phage ELISA) or monoclonal nanobodies (monoclonal ELISA) were added to the wells. After 1 h incubation at 37 °C, the wells were rinsed five times with TPBS. Bound phages were detected by HRP-conjugated anti-M13 antibody (Sino Biological, China), bound VHH were detected by HRP-conjugated anti-Myc-tag antibody (Abcam, UK) followed by the addition of 3,3’5,5’-tetramethylbenzidine (TMB) (Bio-Rad, USA) as a substrate. The reaction was stopped by 1M H_2_SO_4_, the absorbance at 450 nm was read using a Varioskan LUX (Thermo Fisher Scientific, USA).

To confirm RBD-specific binding of the selected nanobodies immunoplates (Nunc MaxiSorp, Thermo Fisher Scientific, USA) were coated with RBD (100 ng per well). Serial two-fold dilutions of antibodies were made, starting from 10 μg/ml to 0.6 ng/ml. 100 μl of each concentration was added to the wells. Bound VHHs were detected by HRP-conjugated anti-Myc-tag antibody and TMB substrate. EC50 values were calculated using four-parameter logistic regression using GraphPad Prism 9 (GraphPad Software Inc, USA).

For competitive ELISA recombinant hACE2 protein were immobilized (400 ng per well) on immunoplates (Nunc MaxiSorp, Thermo Fisher Scientific, USA). RBD-specific nanobodies were mixed with HRP-conjugated recombinant RBD protein (0.2 μg/ml) to two final concentrations of antibodies (0.5 μg/ml and 0.1 μg/ml) and incubated at 37 °C for 30 min. Nanobody to C. difficile Toxin B was used as a negative control. After incubation samples (100 μl) were transferred to hACE2 coated plates and incubated at 37 °C for 30 min. The TMB solution was added for 15 min, the reaction was stopped by the addition of 1M H_2_SO_4_. The inhibition rate of ACE2-RBD binding was calculated by comparing the signals in the sample and the negative control wells.

### Expression and purification of nanobodies

Phagmids encoding the sequences of the selected nanobodies were transformed into BL21 cells (NEB, USA) for expression and purification. The cells were cultured in 100 ml 2xYT (containing 100 μg/ml ampicillin) overnight at 30 °C. The cells were harvested by centrifugation and lysed by a BugBuster Protein Extraction Reagent (Novagen, USA). Nanobodies were purified by IMAC using an AKTA start system and a HisTrap column, dialyzed against PBS and sterile filtered. Protein expression and purity was confirmed with SDS-PAGE.

### Microneutralization assay with live SARS-CoV-2

Nanobodies were serial two-fold diluted starting from 20 μg/ml to 1.2 ng/ml in complete Dulbecco’s modified Eagle medium (DMEM) supplemented with 2% heat-inactivated fetal bovine serum (HI-FBS). Triplicates of each dilution were mixed with 100 50% tissue culture infectious doses (TCID50) of SARS-CoV-2 and incubated at 37 °C for 1 h. After incubation samples were added to a monolayer of Vero E6 cells and incubated in a 5% CO_2_ incubator at 37 °C for 96-120 h. The cytopathic effect (CPE) of the virus was assessed visually. The minimal neutralizing concentration was defined as the lowest antibody concentration (highest antibody dilution) that completely inhibited CPE of the virus in two or three of the three replicable wells.

The following SARS-CoV-2 strains were used in the assay: B.1.1.1, B.1.1.7, B.1.351, B.1.1.28/P.1, B.1.617.2 and B.1.1.529.

All experiments were performed in a Biosafety Level 3 facility (BSL-3).

### Binding kinetics measurements

The affinity of nanobodies was measured by surface plasmon resonance (Biacore 3000, GE Healthcare Bio-Sciences AB, Sweden). Recombinant RBD protein at a concentration 20 μg/ml in 10 mM acetate buffer (pH 4.5) was immobilized on the surface of a CM5 sensor chip using a Amine Coupling Kit (GE Healthcare Bio-Sciences AB, Sweden). VHHs were serial 4-fold diluted from 417 nM to 0 nM and flowed over the captured RBD surface at 15 μl/min for 3 minutes. Dissociation time was 10 minutes. Working buffer was HBS-EP (0.01 M HEPES pH 7.4, 0.15 M NaCl, 2 mM EDTA, 0.005% v/v Surfactant P20). After each injection, the chip surface was regenerated with 10 mM Tris-HCl, pH 1.5 for 40 s at a flow rate 30 μl/min. Calculations were performed using the BIAEvaluation software (GE Healthcare Bio-Sciences AB, Sweden).

### Generation of homodimeric and heterodimeric forms of nanobodies

Homodimeric and heterodimeric forms of nanobodies were generated using two rounds of PCR followed by cloning PCR products into a pHEN1 vector. VHHs in dimeric forms were fused by GS-linker (Gly4Ser)_4_. In the first round of PCR, VHH monomers were amplified using Q5 High-fidelity DNA polymerase (NEB, UK) and the primers listed in Table 2. In the second round PCR nanobodies sequences were assembled together and amplified using pHEN1-F and pHEN1-R primers. PCR products and the pHEN1 vector were digested with SfiI and NotI restriction enzymes (Thermo Fisher Scientific, USA) and ligated using T4 ligase (Thermo Fisher Scientific, USA). Recombinant vectors were used for TG1 cells transformation.

**Table 2.**
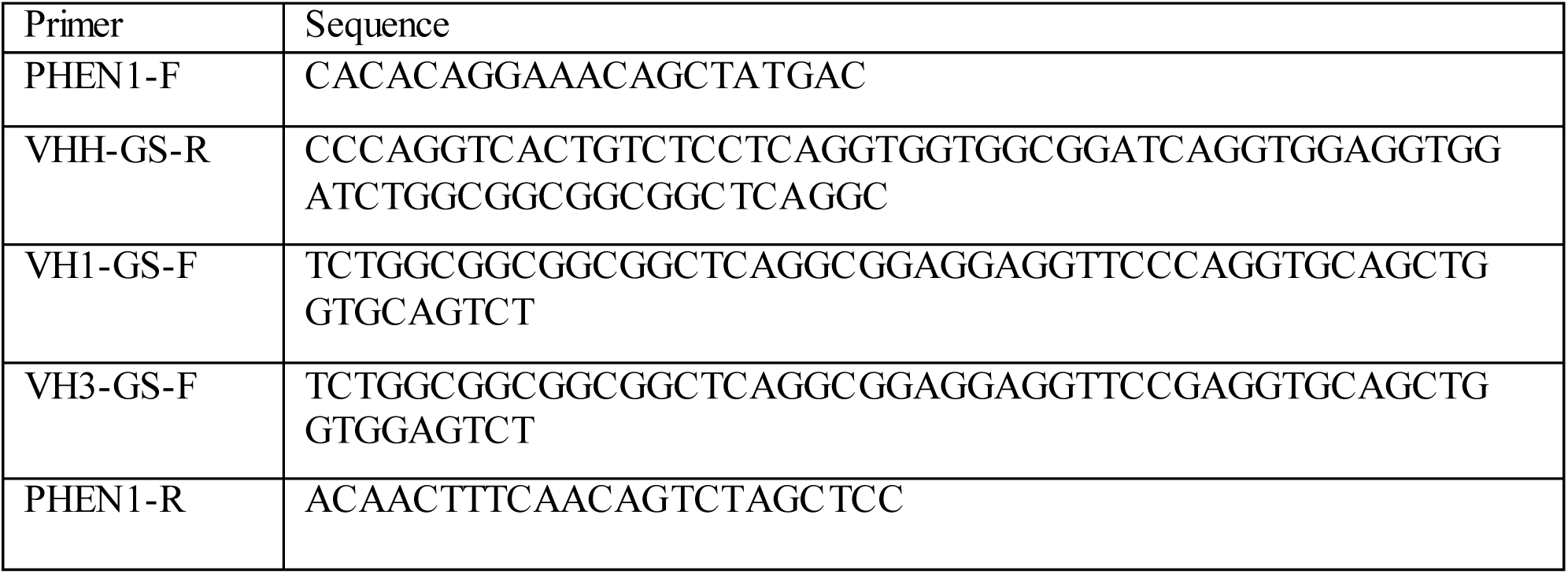
Primers used for generation of dimeric molecules

### DNA isolation and sequencing

Phagemid DNA was isolated from bacterial cells using the Plasmid miniprep kit (Evrogen, Russia). VHH-coding sequences were sequenced with Lac-prom (5’-CTTTATGCTTCCGGCTCGTATG-3’) and pIII-R (5’ CTTTCCAGACGTTAGTAAATG 3’) primers according to the protocol of the BigDyeTerminator 3.1 Cycle Sequencing kit for the Genetic Analyzer 3500 Applied Biosystems (Waltham, MA, USA). The electrophoretic DNA separation was performed in 50-cm capillaries with POP7 polymer. MEGA X was used for the generation of the single consensus sequences by assembling of the forward and reverse sequences (41).

### Phylogenetic tree analysis

A phylogenetic tree of isolated VHHs was constructed to display the sequence diversity of single-domain antibodies. The analysis comprised the following traditional steps: alignment, phylogeny and tree rendering. Sequences were aligned with MUSCLE (v3.8.31) (42). The phylogenetic tree was reconstructed using the maximum likelihood method implemented in the PhyML program (v3.1/3.0 aLRT) (43). The WAG substitution model was selected assuming an estimated proportion of invariant sites (of 0.000) and 4 gamma-distributed rate categories to account for rate heterogeneity across sites. The gamma shape parameter was estimated directly from the data (gamma=0.448). Reliability for internal branch was assessed using the aLRT test (SH-Like) (44). Graphical representation and edition of the phylogenetic tree were performed with MEGA X (41).

## AUTHOR CONTRIBUTIONS

Conception and design of the experiments: I.A.F., D.V.S., I.B.E., I.V.D., library construction, phage display, cloning, production, purification and characterization of VHH and their dimeric forms: I.A.F., I.A.A., A.I.K., I.B.E., E.I.R., A.A.D., O.V.Z., production and purification of recombinant proteins: I.B.E., E.I.R., A.A.D., SARS-CoV-2 culture and microneutralization assay: I.V.D., A.V.K., A.A.I., A.G.B., SRP kinetic measurements: D.V,V., camel immunization and blood collection: D.A.E., DNA sequencing, phylogenetic tree construction: O.L.V., N.N.R., E.I.A., M.S.K., data analysis and interpretation, figures preparation: I.A.F., D.V.S., drafting the manuscript: I.A.F., revision the manuscript: D.V.S., I.B.E., I.V.D., O.L.V., approval of the final version for submission: D.V.S., D.Y.L., B.S.N., A.L.G.

## ACKNOWLEDGMENTS

We would like to thank Y.S. Lebedin who kindly provided us with the recombinant hACE2 protein and the HRP conjugated recombinant RBD protein.

